# Observational epidemiological studies can mitigate genetic confounding with a genetic relatedness matrix

**DOI:** 10.1101/2025.11.24.690292

**Authors:** Roshni A. Patel, Joshua G. Schraiber, Matt Pennell, Michael D. Edge

## Abstract

Observational studies are commonly used in psychology and epidemiology to identify risk factors correlated with health outcomes. However, these studies are vulnerable to confounding when shared genetic variation influences both the putative risk factor and outcome. Researchers have historically controlled for this type of genetic confounding using polygenic scores, but these scores are often noisy and biased estimators of a trait’s genetic component. Here, we develop a method that leverages a genetic relatedness matrix (GRM) to control genetic confounding when testing for non-genetic risk factors. In simulations, we find that our method outperforms existing approaches, particularly at sample sizes that are large by the standards of much human research but smaller than datasets often used in human genetics. We also demonstrate that existing methods are susceptible to poor GWAS portability, whereas our method is inherently robust to such concerns. Finally, we apply our method to the UK Biobank to re-analyze social risk factors for health outcomes in previously understudied cohorts.

**Significance statement:** Large observational datasets are often used in epidemiology or social sciences to identify risk factors associated with health outcomes. However, these studies can be misleading when the putative risk factor and outcome are both complex genetic traits: a correlated genetic basis can create spurious associations between a putative risk factor and outcome. Existing approaches to address this problem typically rely on GWAS summary statistics, which require biobank-scale datasets. In this work, we introduce an alternative method that uses a genetic relatedness matrix (GRM) to control for genetic confounding directly. We show that our approach is closely related to existing methods but outperforms them in smaller datasets, making it an especially valuable tool for researchers who work on understudied traits and populations.

## Introduction

Empirical research across the social, behavioral, and health sciences often relies on observational studies to test for relationships between putative exposures and outcomes. However, these observational studies can be subject to confounding when the exposure and outcome are both complex traits with a genetic component. Complex traits are influenced by many genetic variants, with estimates for some traits reaching into the tens or even hundreds of thousands (Bulik-Sullivan *et al*., 2015; Sinnott-Armstrong *et al*., 2021; Yengo *et al*., 2022). These variants also tend to be pleiotropic, influencing multiple traits simultaneously, which raises the possibility that both exposure and outcome are influenced by shared genetic variation (Bulik-Sullivan *et al*., 2015).

Shared genetic variation poses a problem for causal inference in observational epidemiological studies. In many instances, researchers are interested in understanding whether intervening on a particular exposure (e.g. BMI) could modify someone’s risk for a particular outcome (e.g. high cholesterol). In other words, to understand the effect of environmental interventions, researchers seek to understand whether an observed correlation between exposure and outcome manifests independently of genetics: correlations that arise solely due to shared genetic variation may not readily translate to effective interventions. For example, previous work has found that associations between developmental risk factors and later-in-life psychopathology are attenuated after accounting for shared genetics and other factors (D’Onofrio *et al*., 2014). Consequently, there is a growing awareness of the importance of controlling for shared genetic variation that affects the putative exposure and outcome of interest, both in studies of population samples, as we focus on here, as well as in twin studies (D’Onofrio *et al*., 2020; Eney *et al*., 2017; Jaffee and Price, 2012; Moshtael *et al*., 2024). Within the social sciences, recognition of this problem has motivated a surge of interest in behavioral genetics and the use of genetic information to better understand the effects of behavioral interventions (Harden and Koellinger, 2020).

Historically, efforts to control genetic confounding—specifically, the genetic confounding that results from genetic correlation between exposure and outcome—have relied on including covariates that capture the genetic variation underlying the exposure. One approach conditions on a small number of genetic variants that have a large effect on the exposure trait (Choi *et al*., 2019; Manousaki *et al*., 2021; Davey Smith and Ebrahim, 2003; Davey Smith and Hemani, 2014). However, this approach relies on the genetic architecture of the exposure trait being dominated by common, large-effect variants, which is not always true for complex traits (Sinnott-Armstrong *et al*., 2021). Another line of work has sought to control for genetic confounding by including the polygenic score for the exposure as a covariate when regressing outcome on exposure (Belsky *et al*., 2016; Conley, 2016; Cuevas *et al*., 2021; D’Onofrio *et al*., 2020; Lee *et al*., 2021; Schmitz and Conley, 2017; Zheng *et al*., 2023). Polygenic scores (PGS) estimate the genetic component of a trait by combining information across hundreds or thousands of genetic variants ascertained from genome-wide association studies (GWAS). In theory, one could control genetic confounding by perfectly estimating the true value of the genetic component of the exposure, and including it as a covariate when regressing outcome on exposure. But in practice, PGS estimated from data do not capture all causal variants, and the effects of these causal variants are estimated with bias and noise. In response, a considerable body of work has sought to improve control of genetic confounding by improving PGS estimation and carefully modeling measurement error (DiPrete *et al*., 2018; Pingault *et al*., 2021, 2022; Uddin *et al*., 2022). However, less progress has been made in improving the “portability” of PGS. Due to population structure confounding and challenges with fine-mapping causal variants, PGS are known to be less accurate when estimated from and applied to groups with different genetic ancestries or environments (Martin *et al*., 2019; Mostafavi *et al*., 2020; Wang *et al*., 2024). As a result, PGS-corrected estimates of the exposure effect are likely to be unreliable when applied to populations that are underrepresented in the cohorts from which that PGS was estimated.

Recently, Zhao *et al*. (2024) introduced an alternative approach to control genetic confounding named PENGUIN. PENGUIN-corrected estimates of the exposure effect are a tremendous improvement over previous methods that rely on polygenic scores. Whereas earlier PGS-based methods sought to control genetic confounding by noisily estimating individual-level genetic values, PENGUIN uses LD score regression (LDSC) (Bulik-Sullivan *et al*., 2015) to directly estimate the genetic correlation between exposure and outcome from GWAS summary statistics. This enables PENGUIN to estimate the exposure effect by subtracting off the correlation between exposure and outcome that is mediated through genetics.

However, at least two problems remain. First, both PENGUIN and PGS-based methods presuppose that data on exposure and outcome have been collected in samples large enough to allow very precise GWAS estimates (in practice, often hundreds of thousands of individuals). This requirement limits the utility of existing datasets where valuable phenotypic information may have been collected in smaller sample sizes. Moreover, biobank-scale measurements may be fundamentally untenable for many variables analyzed in observational epidemiological studies: psychologists commonly study traits that are time-consuming and labor-intensive to measure, such as longitudinal phenotypes or scales computed from multi-question surveys. Second, even when traits have been measured in large cohorts, methods that rely on GWAS summary statistics are susceptible to poor portability across cohorts with differences in genetic ancestries or environmental backgrounds (Mostafavi *et al*., 2020; Patel *et al*., 2025; Wang *et al*., 2024). Despite growing recognition of this problem, progress in expanding the diversity of biobank participants remains slow (Mills and Rahal, 2019). Thus, there remains a need to develop a method that controls genetic confounding when studying traits that are hard to measure, or populations underrepresented in existing GWAS.

To develop a flexible method to control genetic confounding in observational epidemiological studies, we propose using a genetic relatedness matrix (GRM). GRMs are typically constructed as variance-covariance matrices of standardized, additive genotypes, and they encode information on genetic relationships among individuals (Speed and Balding, 2015). Thus, the GRM can be used to partition the naive correlation between exposure and outcome into a component mediated through genetics and a component independent of genetics (Lee *et al*., 2012; Yang *et al*., 2010). Here, we demonstrate the conceptual similarity between GRM-corrected estimates and estimates derived from earlier GWAS-based methods. We show that GRM-corrected estimates largely resolve the two problems associated with GWAS-corrected estimates: they can be applied to small datasets and computed within-sample, mitigating portability concerns. We conclude by using our method to reanalyze social risk factors for health outcomes in 6,104 Indian and 8,483 Black British individuals in the UK Biobank.

## Results

### Statistical model

We begin by introducing a generative statistical model of two phenotypes that we refer to as the exposure, *X*, and the outcome, *Y* (Figure 1A):

**Figure 1.**
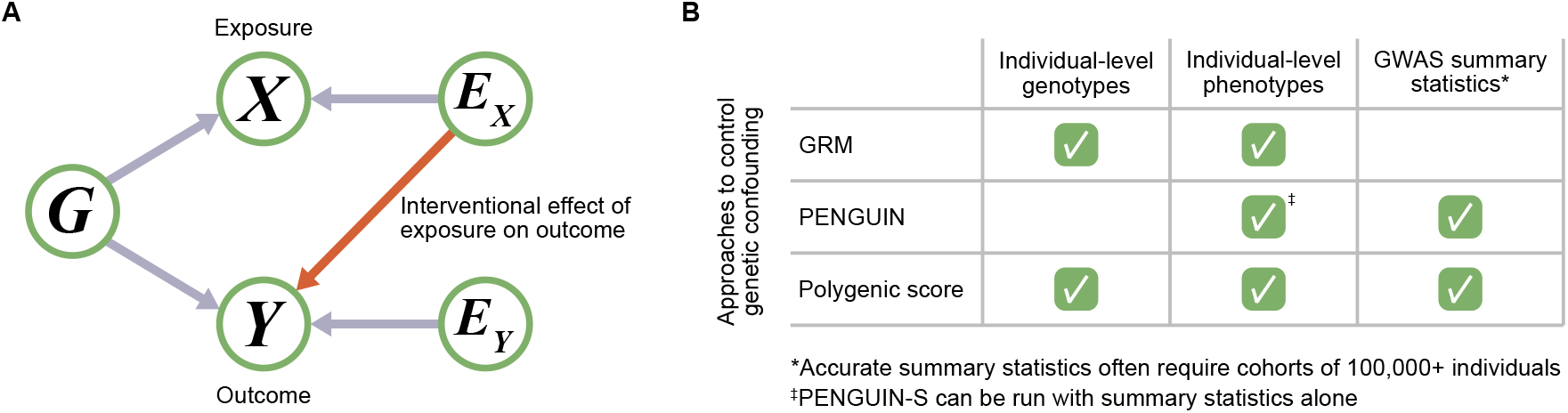
Overview. **A)** Causal graph for observational epidemiological studies where both exposure and outcome are genetic traits. The orange arrow illustrates the quantity of interest: the interventional effect of the exposure on the outcome. **B)** Data requirements for GRM-, PENGUIN-, and PGS-corrected estimates of exposure effect.

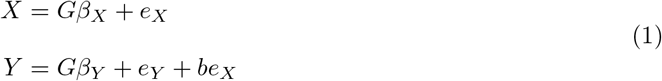

where, for *n* individuals and *m* genetic variants, *G* is the centered *n × m* genotype matrix; *β*_*X*_ and *β*_*Y*_ are *m ×* 1 vectors of additive SNP effect sizes; and *e*_*X*_ and *e*_*Y*_ are *n ×* 1 vectors of non-genetic effects on the traits. We assume the entries of *e*_*X*_ and *e*_*Y*_ are independently and identically distributed across individuals with expectations of 0 and that *e*_*X*_ and *e*_*Y*_ are uncorrelated. We further assume that non-genetic effects *e*_*X*_ and *e*_*Y*_ are independent of *G*. We assume *β*_*X*_ and *β*_*Y*_ have expected values of 0, and we allow that *β*_*X*_ and *β*_*Y*_ could be correlated.

Under this generative model, the outcome is correlated with the exposure through both genetic and non-genetic mechanisms, and the coefficient *b* measures the effect of the non-genetic component of the exposure. In other words, *b* is the quantity we wish to estimate in observational epidemiological studies: it quantifies the extent to which intervening on the exposure will affect the outcome variable. In contrast, the naive estimate of the exposure effect—obtained by simply regressing the outcome on the exposure—is 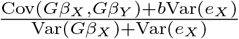 in expectation. In other words, the naive estimate of the exposure effect is subject to both regression attenuation and bias from genetic correlation between traits (see Supplementary Note 1 for details).

Although this model has not been explicitly formalized in prior literature (to our knowledge), we show that it can be understood as the basis for including the exposure PGS as a covariate and other existing approaches to control genetic confounding (Supplementary Note 1). Given that *e*_*X*_ is unobserved and unmeasurable, estimating *b* requires an indirect approach. PENGUIN approaches this problem by modeling Cov(*Gβ*_*X*_, *Gβ*_*Y*_), whereas PGS-based methods seek to estimate *Gβ*_*X*_. Here, we propose estimating the interventional effect of the exposure by using a GRM to partition the covariance between exposure and outcome into a genetic component and a non-genetic component. If we assume that both the effects of genetic variants and the non-genetic effects on traits are normally distributed, the exposure and outcome can be modeled with a bivariate normal distribution:

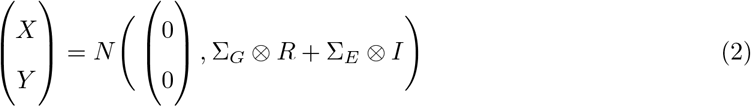

where *R* is an *n × n* GRM, *I* is the *n*-dimensional identity matrix, and Σ_*G*_ and Σ_*E*_ represent the 2 *×* 2 genetic and non-genetic variance-covariance matrices between exposure and outcome. Given a set of genotypes from which to construct the GRM *R*, and phenotypes for the exposure *X* and outcome *Y*, we can estimate the genetic and non-genetic variance components Σ_*G*_ and Σ_*E*_ by using variance component analysis to model the covariance of *X* and *Y* across individuals. In fact, the model in Equation 2 may be familiar as bivariate GREML, which is commonly used to estimate heritability and genetic correlation using the entries of Σ_*G*_ (Yang *et al*., 2010; Lee *et al*., 2012). In contrast, we propose using the entries of Σ_*E*_ to estimate the interventional effect of an exposure in the context of observational epidemiological studies. Specifically, let the entries of Σ_*E*_ be denoted as follows:

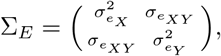

where 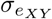 is the covariance between the non-genetic components of the exposure *X* and outcome 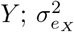 is the variance of the non-genetic component of the exposure *X*; and 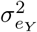 is the variance of the non-genetic component of the exposure *Y*. As implied by equation 1, the covariance between the non-genetic component of the exposure *X* (*e*_*X*_) and the non-genetic component of the outcome 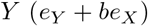 *Y* (*e*_*Y*_ + *be*_*X*_) is 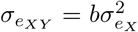. Thus, an estimator for the interventional effect of the exposure *X* is given by:

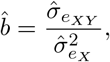

where 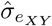 and 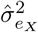 are estimators of the corresponding quantities—in this paper, the ones provided by GREML.

### Assessing the utility of GRM-corrected estimates

Though PGS-, PENGUIN-, and GRM-corrected estimates share the same underlying model of trait variation, these approaches impose different data requirements (Figure 1B). PGS-corrected estimates require GWAS summary statistics to construct a PGS and individual-level genotypes and phenotypes to regress outcome on exposure. In contrast, PENGUIN-corrected estimates do not require individual-level genotypes. Instead, PENGUIN uses GWAS summary statistics to estimate the genetic correlation between traits, and individual-level phenotypes to estimate the overall phenotypic correlation between exposure and outcome. Finally, GRM-corrected estimates require no GWAS summary statistics at all and instead partition the covariance between exposure and outcome strictly using individual-level genotypes and phenotypes.

Thus, we sought to understand the utility of different methods for controlling genetic confounding. We simulated genotypes for individuals under a realistic model of human demography (Gutenkunst *et al*., 2009; Adrion *et al*., 2020). Conditional on these genotypes, we simulated phenotypes for exposure and outcome traits under the generative model described in Equation 1 and an additive genetic architecture with frequency-dependent effect sizes (see Methods).

In simulations, we find that the perfect PGS—that is, the true value of the genetic component of the exposure—would control genetic confounding so successfully that the resulting estimates lie on the diagonal *y* = *x* line (Figure 2A). However, as the correlation between the estimated PGS and the true genetic value decreases, PGS-corrected estimates of the exposure effect become increasingly susceptible to genetic confounding. Any genetic effects not captured by the estimated PGS are absorbed into the coefficient of the exposure, which is why the resulting estimate of exposure effect is biased towards the genetic correlation between exposure and outcome. Strikingly, even when a PGS is constructed with 100% of the true causal variants, noisy estimation of variant effect sizes will result in biased estimates of exposure effect. Given that current GWAS are not expected to identify 100% of the causal variants for a trait, this represents a relatively optimistic scenario. When only 20% of causal variants are captured, the PGS-corrected estimate of exposure effect barely performs better than the estimate from the naive regression of outcome on exposure. For the remainder of this manuscript, we therefore prioritize comparing GRM-corrected estimates with PENGUIN-corrected estimates.

**Figure 2.**
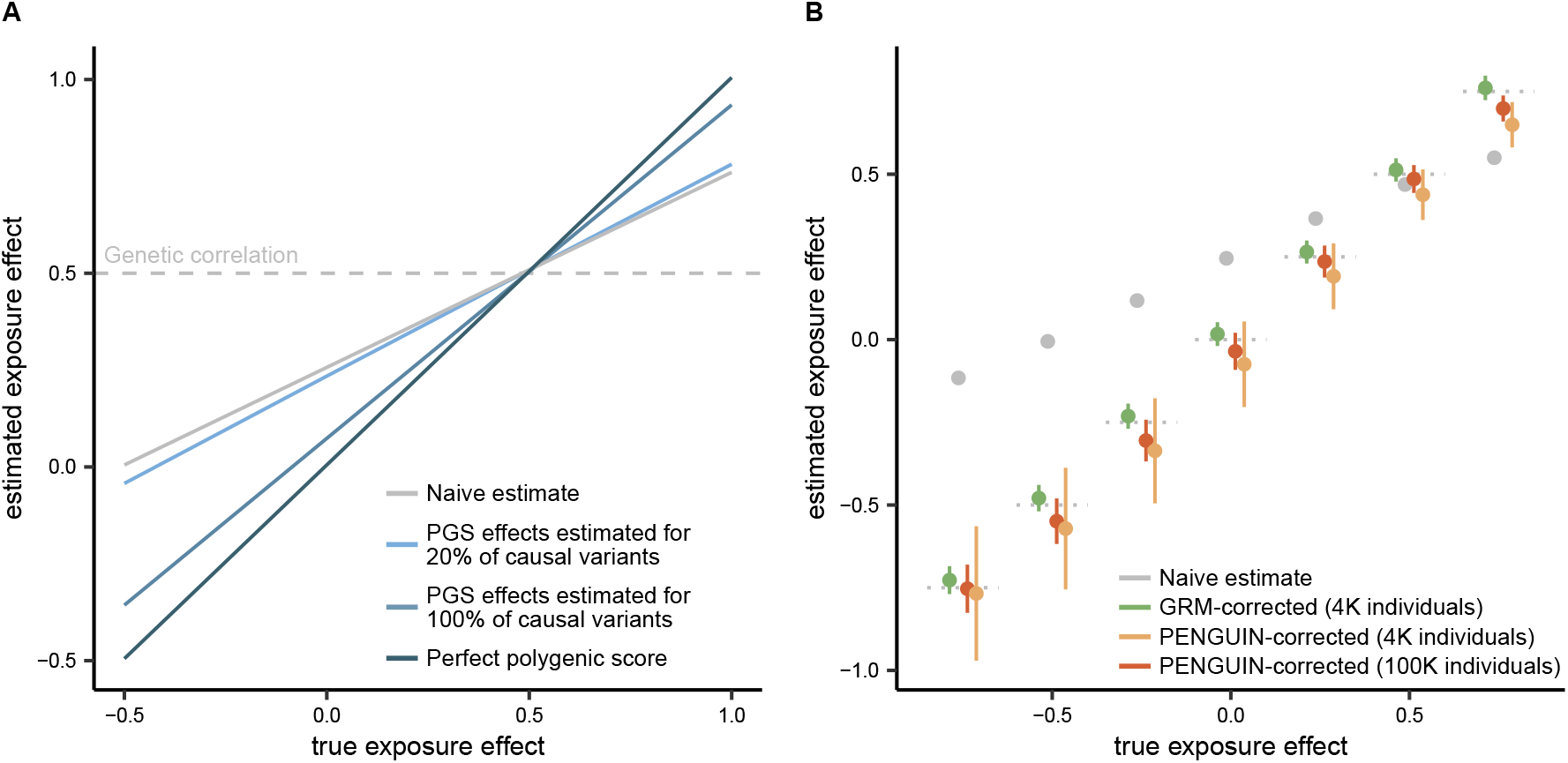
Utility of GRM-corrected estimates. **A)** PGS-corrected estimates of exposure effect, for i) PGS constructed from 20% of causal variants and effects estimated with noise; ii) PGS constructed from 100% of causal variants and effects estimated with noise; and iii) the perfect PGS, which exactly overlaps the diagonal, i.e. the *y* = *x* line. **B)** GRM- and PENGUIN-corrected estimates of exposure effect in simulated data. Horizontal dotted lines correspond to the diagonal, i.e. the *y* = *x* line. Error bars correspond to mean *±* 1 standard deviation. In both panels, the naive estimate is obtained by directly regressing outcome on exposure.

We next compared the performance of PENGUIN- and GRM-corrected estimates in realistic scenarios (Figure 2B). Some observational epidemiological studies are conducted in biobanks with information on hundreds of thousands of individuals, whereas others rely on datasets with a few thousand individuals (Conley, 2016; D’Onofrio *et al*., 2020). We find that GRM-corrected estimates derived from sample sizes of just 4,000 individuals perform as well as PENGUIN-corrected estimates derived from biobank-scale data. In smaller sample sizes, we find that GRM-corrected estimates substantially outperform PENGUIN-corrected estimates, reflecting the reliance of PENGUIN on GWAS summary statistics. At a sample size of 4,000 individuals, GWAS summary statistics are too noisy for accurate estimation of heritability and genetic correlation via LD score regression. This highlights the particular utility of GRM-corrected estimates in studies with sample sizes that are too small for GWAS. On the other hand, the utility of PENGUIN-corrected estimates becomes apparent as sample sizes approach hundreds of thousands of individuals. In this setting, constructing a GRM becomes computationally infeasible, but GWAS summary statistics become less noisy and more accurate.

### Interrogating assumptions about trait heritability

To identify the potential limitations of GRM- and PENGUIN-corrected estimates, we next examined their robustness to assumptions about trait heritability. Both approaches estimate the interventional effect of the exposure by partitioning the genetic correlation between traits from the non-genetic correlation between them, which requires making assumptions about how trait heritability is distributed across individuals and variants. Thus, we sought to understand how violations of these assumptions affect our ability to recover the true interventional value of the exposure effect.

We first evaluated the effect of violating assumptions about random mating. Population structure and cross-trait assortative mating can both induce spurious correlations between individuals’ genotypes and are known to bias traditional heritability and genetic correlation estimators (Border *et al*., 2022, 2024). To characterize the effect of population structure, we simulated two subpopulations with an *F*_*ST*_ of 0.005, a value comparable to those observed between populations within continental Europe. We find that when the mean exposure phenotype is shared between subpopulations, the confidence intervals for GRM- and PENGUIN-corrected estimates generally include the true exposure effect, though PENGUIN-corrected estimates have much wider confidence intervals (Figure S1A). However, if the mean enviromental contributions to the exposure and outcome (that is, the subpopulation-mean values of *e*_*X*_ and *e*_*Y*_) differ between subpopulations (e.g., due to environmental differences), we find that PENGUIN-corrected estimates exhibit greater bias than GRM-corrected estimates (Figure S1B). In simulations of cross-trait assortative mating, we find that confidence intervals for GRM-corrected estimates generally recover the true exposure effect, but estimates can be prone to bias as the strength of cross-trait assortative mating increases (Figure S1C). We were unable to evaluate PENGUIN-corrected estimates under assortative mating given the large sample sizes required for PENGUIN and the computational complexity of assortative mating simulations (Border *et al*., 2024).

Lastly, we considered assumptions about how heritability is distributed across variants. We find that if the exposure and outcome are simulated with a heritability model that violates method assumptions, both GRM- and LDSC-based estimates of heritability will be biased, consistent with previous findings (Supp Fig S2A) (Speed *et al*., 2017). Nevertheless, we find that GRM-corrected estimates of exposure effect are typically more successful at recovering the true effect (Supp Fig S2B).

### Portability and methods to control genetic confounding

Finally, we sought to characterize the portability of methods to control genetic confounding. Whereas GRM-corrected estimates can only be obtained from a single dataset with individual-level genotype and phenotype data, both PGS- and PENGUIN-corrected estimates can be obtained by combining information across multiple cohorts. In the case of PENGUIN-corrected estimates, this feature presents a considerable methodological advantage: by leveraging external GWAS summary statistics, researchers can use PENGUIN to estimate exposure effect even in the absence of individual-level genotype data (Figure 1B). At the same time, combining information across cohorts makes both PENGUIN- and PGS-corrected estimates susceptible to concerns about poor portability across cohorts.

To characterize the portability of PGS- and PENGUIN-corrected estimates, we first distinguish between the types of portability concerns relevant to both methods (Figure 3A). The portability of PGS has been written about extensively (Berg *et al*., 2019; Martin *et al*., 2019; Mostafavi *et al*., 2020; Sohail *et al*., 2019; Wang *et al*., 2024), and is defined as a loss of accuracy when GWAS are conducted in one cohort and the resulting PGS is applied to a second cohort. This loss of accuracy has many causes, including differences in allele frequency of causal variants, differences in LD between causal variants and tag variants identified in GWAS, differences in the role of the environment, and gene-by-gene or gene-by-environment interactions (Mostafavi *et al*., 2020; Patel *et al*., 2022, 2025; Wang *et al*., 2020; Yair and Coop, 2022).

**Figure 3.**
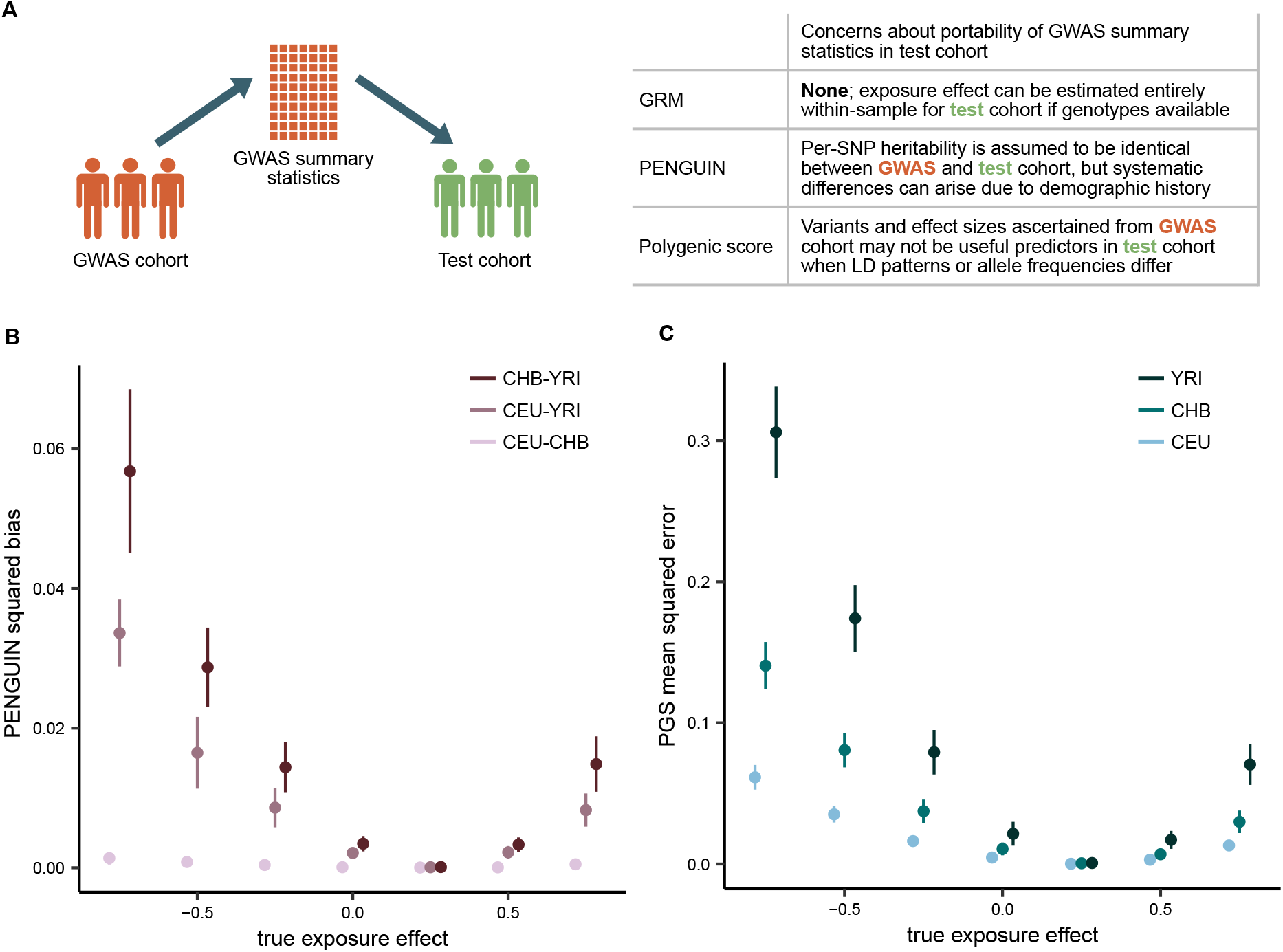
Portability of methods. **A)** Overview of portability concerns associated with GRM-, PENGUIN-, and PGS-corrected estimates of exposure effect. **B)** Squared bias in PENGUIN-corrected estimates of exposure effect when GWAS summary statistics and individual-level data are obtained from two different populations. **C)** Mean squared error in PGS-corrected estimates of exposure effect when GWAS summary statistics are obtained from simulated CEU individuals. Error bars correspond to mean *±* 1 standard deviation.

In contrast, the PENGUIN estimator faces a different kind of portability challenge. In the context of the PENGUIN estimator, “poor portability” means that PENGUIN-corrected estimates are inaccurate when GWAS summary statistics and individual-level phenotypes are obtained from two different cohorts. We show analytically that the PENGUIN estimator is no longer guaranteed to recover the true exposure effect when per-SNP heritability differs between the two cohorts due to factors such as demographic history or selection shaping allele frequencies, or cohort-specific gene-by-gene and gene-by-environment interactions modifying effect sizes (Supplementary Note 2). Specifically, we find that the expected value of the PENGUIN estimator is

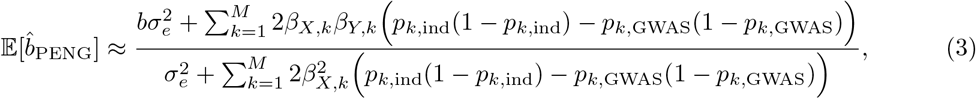

where *β*_*X,k*_ and *β*_*Y,k*_ are the effect of the *k*th variant on the exposure *X* and the outcome *Y*, and *p*_*k*,ind_ and *p*_*k*,GWAS_ are the frequency of the *k*th variant in the individual-level and GWAS cohorts respectively.

To clarify the magnitude of error associated with these two types of portability concerns, we simulated populations under a previously published demographic model inferred from 1000 Genomes CEU (Utah residents with Northern and Western European ancestry), CHB (Han Chinese in Beijing, China), and YRI (Yoruba in Ibadan, Nigeria). Importantly, the portability concerns we modeled are purely due to differences in allele frequency and linkage disequilibrium arising from demographic history. We do not consider gene-by-environment interactions, population structure, or population-specific exposure effects, all of which could further reduce the portability of methods to control genetic confounding.

To characterize PENGUIN portability, we computed the squared bias of the estimator as 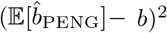. As before, we find that the squared bias increases as the true exposure effect becomes more dissimilar to the genetic correlation (Figure 3B). We also find that the error is most pronounced for comparisons between CEU and YRI, or CHB and YRI. This reflects systematically different per-SNP heritabilities in CEU and CHB relative to YRI, and can be explained by the different amounts of genetic drift in their respective demographic histories.

We next characterized PGS portability by comparing the mean squared error associated with out-of-sample GWAS and in-sample GWAS. Briefly, we performed GWAS on the exposure trait in simulated CEU individuals, clumped trait-associated variants to construct a PGS, and applied that PGS to control genetic confounding in CHB and YRI. Similar to PENGUIN, we find that PGS-corrected estimates exhibit poor portability between groups (Figure 3C). However, we reiterate that unlike PENGUIN, PGS-corrected estimates can be biased even with in-sample GWAS (Figure 2A).

### Empirical data analyses

Having demonstrated the utility of the GRM in controlling genetic confounding, as well as its robustness to assumptions about trait heritability, we turned our attention to empirical data. Specifically, we sought to understand the effect of loneliness on health outcomes in the UK Biobank. Chronic loneliness has an incidence of 5-10% in modern societies (Beutel *et al*., 2017; Hakulinen *et al*., 2018), and is recognized as a significant public health concern (Office of the Surgeon General, 2023). Using observational epidemiological studies, researchers have found that loneliness is associated with numerous health and social outcomes, including increased risk for cardiovascular disease (Hakulinen *et al*., 2018; Liu *et al*., 2024; Paul *et al*., 2021; Rødevand *et al*., 2021; Xia and Li, 2018), increased inflammation (Matthews *et al*., 2024; Van Bogart *et al*., 2022), and poor mental health (Abdellaoui *et al*., 2019; Dennis *et al*., 2021; Lee *et al*., 2021; Rødevand *et al*., 2021). At the same time, loneliness seems to be genetically correlated with some of these same health outcomes (Abdellaoui *et al*., 2019; Moshtael *et al*., 2024), raising the possibility that it could be useful to control for potential genetic confounding when testing for loneliness as a risk factor.

We estimated the effect of a binary loneliness variable on three cardiovascular traits, four mental health traits, and two inflammation markers in the UK Biobank. Given that risk factors can manifest differently across social and environmental contexts (Nianogo *et al*., 2022; Proto and Quintana-Domeque, 2021; Williams *et al*., 2012), we separately analyzed the effect of loneliness in three subsets of the UK Biobank: a random sample of 7,000 individuals who identified as White and British (here-after “White British”), 6,104 individuals who identified as Indian, and 8,483 individuals who identified as Black or Black British (hereafter “Black British”). For each subset, we computed both GRM- and PENGUIN-corrected estimates of the effect of loneliness. To compute PENGUIN-corrected estimates, we used GWAS summary statistics generated from 361,194 White British individuals.

Across all three groups, we generally find that loneliness either has no effect or is associated with poorer health outcomes, corroborating existing literature (Figure 4). We find that GRM- and PENGUIN-corrected estimates are most concordant for White British individuals (*r*^2^ = 0.96, compared with 0.85 for Black British and 0.92 for Indian). This likely reflects the fact that PENGUIN-corrected estimates face portability challenges when individual-level phenotypes and GWAS summary statistics are obtained from two different populations.

**Figure 4.**
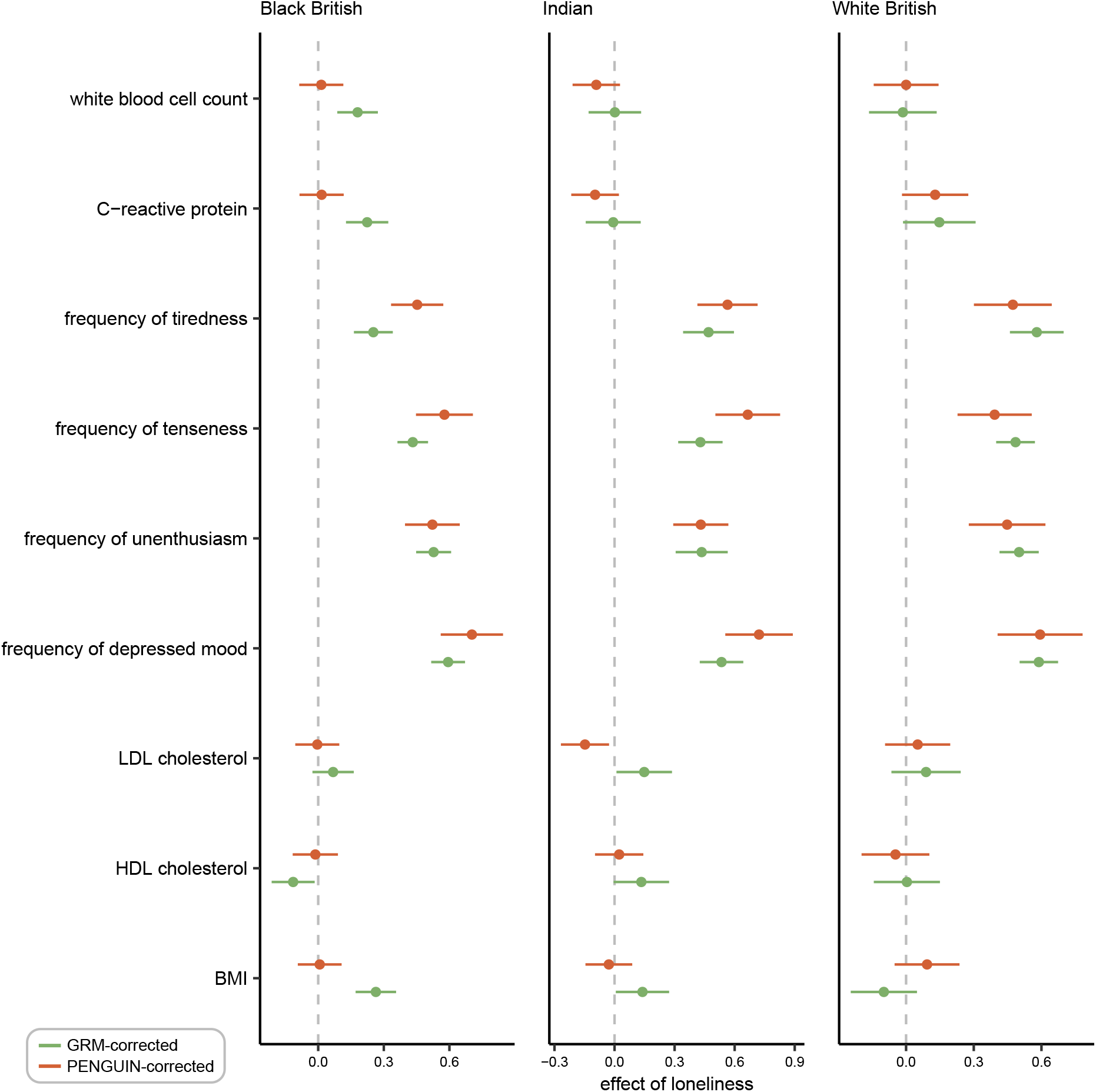
Effect of loneliness on health outcomes in UK Biobank. GRM- and PENGUIN- corrected estimates of the effect of loneliness on inflammatory markers, mental health traits, and cardiovascular risk factors in the UK Biobank Black British, Indian, and White British. PENGUIN-corrected estimates use individual-level data from the specified group and GWAS summary statistics from the UK Biobank White British. Error bars correspond to mean *±* 1 standard deviation. Quantitative traits were inverse normal transformed, such that the x-axis scale has the same interpretation for all quantitative traits, and a different interpretation for categorical mood frequency phenotypes.

This has implications for interpreting the effect of loneliness on health outcomes. For example, PENGUIN trained on a White British GWAS finds no effect of loneliness on inflammation markers in Black British individuals. However, GRM-corrected estimates indicate that loneliness is associated with both increased white blood cell count and increased C-reactive protein. This is particularly notable because loneliness does not appear to have any effect on inflammation in the Indian or White British subsets – thus, the association between loneliness and inflammation is only evident with GRM-corrected estimates, and only in the Black British subset. In some rare instances, we also find that PENGUIN-corrected estimates and GRM-corrected estimates have opposite signs, though we caution that these differences are not significant.

## Discussion

We presented an approach that uses the genetic relatedness matrix (GRM) to control genetic confounding in observational epidemiological studies. Echoing earlier work (Bulik-Sullivan, 2015), our results reiterate the conceptual similarity between LDSC- and GRM-based estimates of heritability. We show that the estimator obtained by the LDSC-based method PENGUIN is equivalent to the estimator obtained when the GRM is used to control genetic confounding. In parallel, our work also highlights the deep connections between statistical genetics and phylogenetics (Schraiber *et al*., 2024). When studying ecological and evolutionary relationships between traits, phylogeneticists often use interspecific data to test for an association between two traits. Crucially, researchers must account for confounding that results from species’ shared ancestry via their phylogenetic relationships (Felsenstein, 1985; Lynch, 1991; Martins and Hansen, 1997; Uyeda *et al*., 2018). This problem in phylogenetics bears a strong resemblance to genetic confounding in observational epidemiological studies, and in fact, the model we propose is closely analogous to one recently described by Westoby *et al*. (2023) in the context of controlling phylogenetic confounding.

Our results provide several lessons for empirical studies. First, our results showcase the value of the GRM in settings where large, accurate GWAS are not available for a given trait or population. When biobank-scale data or GWAS summary statistics for the specific trait and population are readily available, existing methods such as PENGUIN are well-suited to the task. For understudied traits or populations, however, in-sample GWAS will be too noisy for accurate estimation, and out-of-sample GWAS risk portability concerns. In this setting, observational epidemiological studies can benefit from using GRM-corrected estimates of exposure effect.

Second, our results highlight the importance of cohort-specific analyses in epidemiological research aimed at identifying effective interventions. In the UK Biobank, we observed differences in the effect of loneliness on multiple health and social outcomes across Black British, Indian, and White British individuals. The differences we observe could represent either gene-by-environment interactions, environment-by-environment interactions, or both, given that both genetic and non-genetic factors could vary among groups. Leveraging the GRM to characterize such interactions is an important direction for future research.

One important consideration in applying any data-analytic approach is computational tractability. For the results presented here, we performed variance-components analysis using GCTA-GREML, a method that partitions trait variance using restricted maximum likelihood. GCTA-GREML and similar methods become increasingly slow and computationally expensive as sample sizes increase into the tens of thousands, largely because of the difficulty of forming and storing the GRM, which has *n*^2^ entries for a sample of size *n*. In this setting, a promising alternative is SCORE, a recent method developed by Wu *et al*. (2022). Much like GCTA-GREML, SCORE incorporates individual-level genotype data to partition variance for two traits. Unlike GCTA-GREML, however, SCORE uses stochastic trace estimation to avoid explicitly forming a GRM, allowing it to scale to hundreds of thousands of individuals. Although we do not evaluate SCORE in this study, it appears well suited to sample sizes that are too small for GWAS yet too large for GCTA-GREML.

Another important consideration is the construction of the GRM, which is central to our approach. There are different ways to construct GRMs, corresponding to different assumptions about how genetic similarity between individuals translates into phenotypic similarity. In this work, we use what is sometimes called the “canonical” GRM, defined as the variance–covariance matrix of additive genotype values that are standardized by the estimated standard deviation of the genotype assuming Hardy–Weinberg equilibrium. Although widely used, this construction is not necessarily optimal: it measures similarity at observed variants, which may differ from genome-wide relatedness, and it corresponds to an implicit assumption that 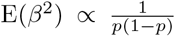, such that the squared effect sizes of variants are inversely proportional to allelic heterozygosity. In our simulations, the canonical GRM generally provides adequate control of genetic confounding, even when the data are generated under alternative models. Nevertheless, alternative GRMs could readily be incorporated into observational epidemiological studies. In particular, recent work has highlighted the advantages of GRMs computed on the basis of estimated ancestral recombination graphs, especially when not all variants are genotyped (Fan *et al*., 2022; Link *et al*., 2023; Zhang *et al*., 2023; Lehmann *et al*., 2025). Other alternatives include GRMs that encode different assumptions about the effect size-allele frequency relationship, or explicitly incorporate information on linkage disequilibrium (LD) or heritability (Speed *et al*., 2017). The performance of the GRM under different generative models also remains to be characterized: although our simulations incorporated several complexities, including population structure and assortative mating, many alternative data-generating mechanisms remain unexplored.

Our results present an opportunity to reconsider the causal model underlying observational epidemiological studies. In this work, we focused on estimating the effect of the exposure under a specific causal model where potentially correlated genetic variation affects both exposure and outcome, and researchers are interested in the interventional effect of the exposure, i.e. the effect of the non-genetic component (Fig. 1A). We show in Supplementary Note 1 that this model underlies a large body of existing work in epidemiology and social science that seeks to control genetic confounding by using the polygenic score for the exposure. Nevertheless, we will briefly discuss several alternative causal models that might occur to readers (Figure 5). Two of these models, though apparently different from the generative model displayed in Figure 1, are in fact special cases of it, and estimates derived from our approach (or, given the appropriate conditions, from PENGUIN) will provide germane information about causal relationships. However, the third is an extension of the model that requires a different interpretation.

**Figure 5.**
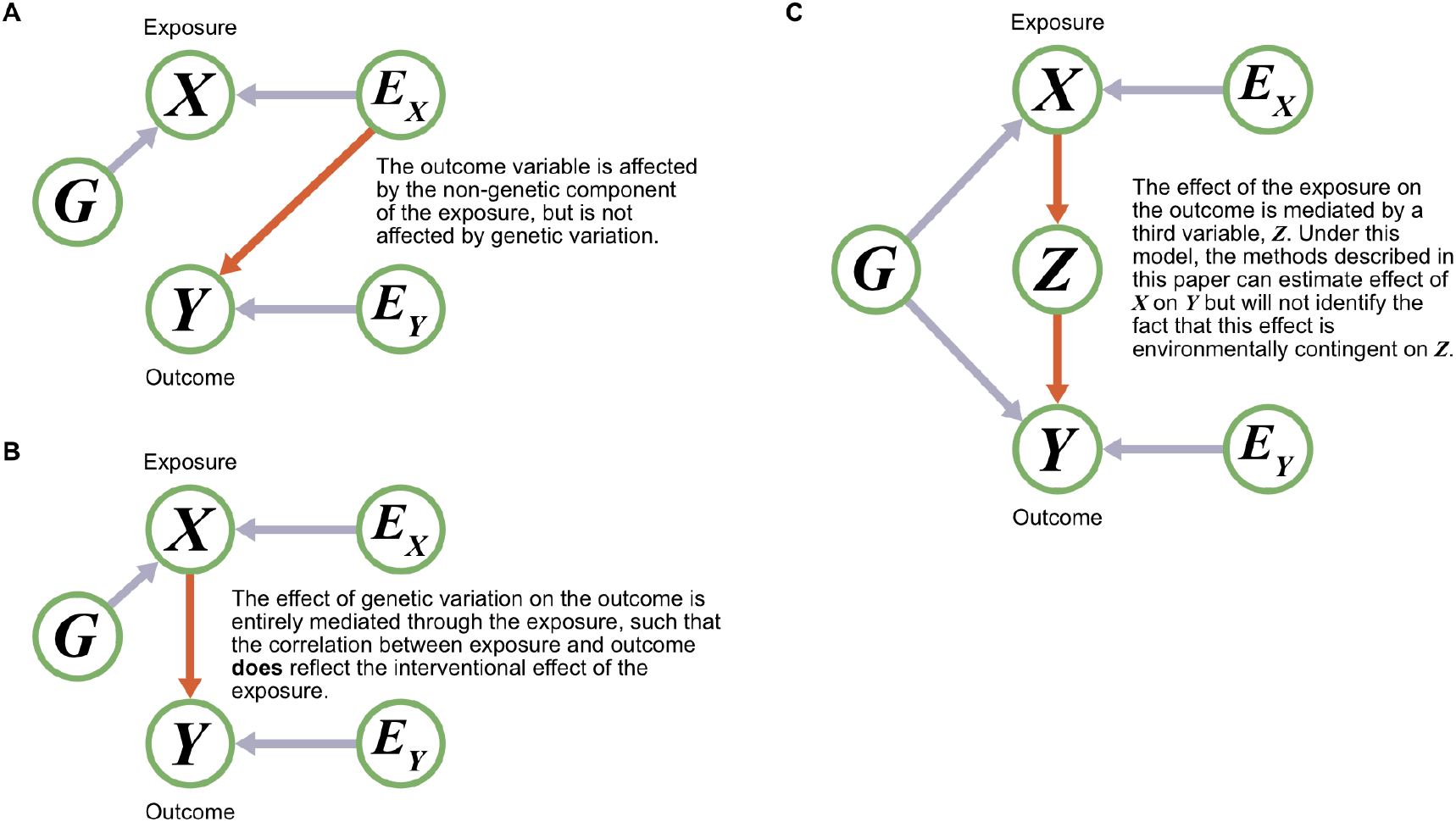
Alternative causal models. **A)** There is no effect of genetic variation on the outcome trait. **B)** The effect of genetic variation is entirely mediated through the exposure trait. **C)** The effect of genetic variation is mediated through the exposure trait, and the effect of the exposure is itself mediated by a third environmental variable.

First, suppose that there is no effect of genetic variation on the outcome trait (Figure 5A). This can be viewed a special case of the generative model in Figure 1A in which *β*_*Y*_ = 0, and therefore the genetic correlation between exposure and outcome is 0. In this setting, the naive regression of outcome on exposure is not subject to genetic confounding, but as we see in Supplementary Note 1, the naive estimator is still subject to regression attenuation. This motivates the use of GRM- or PENGUIN-corrected estimates of exposure effect even when researchers believe the causal model at hand is Figure 5A. In principle, both methods should successfully estimate the exposure effect even if the heritability of the outcome trait is 0.

A second possible scenario is one in which the effect of genetic variation on the outcome is entirely mediated through the exposure (Figure 5B). In this setting, it seems plausible that researchers may want to operationalize a different definition of “interventional effect.” In keeping with the existing literature, our original model defines the interventional effect of the exposure as the effect of the non-genetic component of the exposure. In contrast, in Figure 5B, it may appear more suitable to consider the interventional effect as the effect of the total exposure on the outcome, rather than simply the effect of the non-genetic component of the exposure. However, Figure 5B can be viewed as a special case of our generative model where exposure and outcome are perfectly genetically correlated, i.e. *β*_*Y*_ = *cβ*_*X*_ for some scalar *c*. In this setting, it turns out that the naive regression of outcome on exposure is statistically unbiased and equal in expectation to GRM- and PENGUIN-corrected estimates.

In contrast to the models in Figure 5A-B, the model in Figure 5C is not a special case of generative model in Figure 1A. In particular, now the effect of the exposure on the outcome is mediated by another variable, *Z*, which might be environmentally contingent. This model is motivated by Jencks’ famous “red-headed child” thought experiment, which he pithily summarized, “If, for example, a nation refuses to send children with red hair to school, the genes that cause red hair can be said to lower reading scores. This does not tell us that children with red hair cannot learn to read” (Jencks *et al*., 1972). In Jencks’ example, there is a “genetic” effect of red hair on educational attainment, but it is transmitted via a societal decision to discriminate against red-haired children. A model like ours might indeed suggest, in Jencks’ world, that an environmental modification of hair color— perhaps via sun exposure, bleach, or dye—might affect a child’s educational attainment. But the data analysis suggested here would not, on its own, reveal the environmental contingency of the effect of red hair on educational attainment, which is the crucial consideration in assessing its modifiability. We raise this case to note that the ability to draw causal conclusions from variance decompositions rests on simplifying and sometimes restrictive assumptions (Lewontin, 1974). Thus, the outputs of our approach, despite their favorable properties under the model assumed here and in previous work, should always be treated with caution.

Ultimately, our results position the GRM as a flexible tool for addressing genetic confounding in observational epidemiological studies, particularly in settings where large, well-powered GWAS are unavailable. More broadly, our findings underscore the methodological similarities between disparate lines of work in epidemiology, statistical genetics, and phylogenetics—and the ways in which these connections can be leveraged to improve estimation across domains.

## Materials and methods

### Estimating exposure effect

#### GRM-corrected estimates

To construct a GRM for a particular set of individuals, we used all biallelic variants with minor allele frequency greater than 0.01. Unless specified otherwise, the GRM was constructed with an algorithm corresponding to the assumption that all variants contribute equally to heritability (GCTA flag --make-grm-alg 0).

To compute GRM-corrected estimates of the interventional exposure effect, we used bivariate GREML (Lee *et al*., 2012) to decompose the variance of the exposure and outcome into genetic and non-genetic components. We estimated the exposure effect using the estimated variance-covariance matrix for the non-genetic components of the exposure and outcome. Specifically, we estimated *b* as

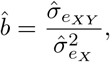

where 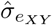 is the estimated covariance between the non-genetic components of the exposure and outcome, and 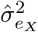 is the estimated variance of the non-genetic component of the exposure.

To obtain standard errors of the GRM-corrected estimator, we used two approaches. First, we computed the standard deviation of the estimates obtained across multiple replicates in simulations. Second, we approximated the standard error of the estimator using the delta method:

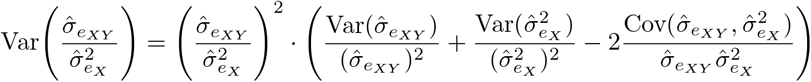

To obtain the variance and covariance of the estimated components, we lightly modified GCTA (https://github.com/roshnipatel/GCTA) to report the Fisher information matrix, which is already computed by the software during each REML iteration. We verified in simulations that both standard errors were comparable (Supp Fig S3).

#### PENGUIN-corrected estimates

To compute PENGUIN-corrected estimates in simulated data, we first generated GWAS summary statistics for biallelic variants with minor allele frequency greater than 0.01 using plink2 (Chang *et al*., 2015). We next generated LD scores for our simulated genotypes using LDSC (Bulik-Sullivan *et al*., 2015) with a window size of 100 kb. Using our GWAS summary statistics and LD scores, we ran individual-level PENGUIN with default parameters. In simulations of random mating, we did not include additional covariates when running GWAS or PENGUIN. In simulations of population structure or assortative mating, we included one principal component when generating GWAS summary statistics and subsequently computing PENGUIN-corrected estimates of exposure effect.

#### PGS-corrected estimates

Due to the many technical decisions necessary to construct polygenic scores (Lin *et al*., 2025), we sought to place an upper bound on the utility of PGS-corrected estimates through semi-analytical means. We simulated trait architecture and genotypes and phenotypes for 100,000 individuals as described below (see *Random mating*). We then simulated the performance of PGS-corrected estimates in scenarios in which 20% and 100% of causal variants are identified in GWAS and included in the polygenic score. To do so, we started by randomly sampling the aforementioned proportion of causal variants, given that all variants contribute equally to heritability in expectation under the model we assume (see *Random mating*). In particular, for the subset of causal variants identified in GWAS, we simulated effect sizes estimated with noise using a standard approach (Vilhjálmsson *et al*., 2015). Estimated effect sizes 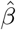 were drawn from a normal distribution centered on the true effect size, with a variance that depends on the true effect size, the allele frequency of the variant, *f*; and the number of individuals in the GWAS, *n*.

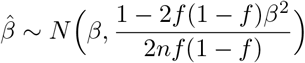

Using the estimated effect sizes 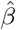, we then simulated an estimated polygenic score for each individual. Using this estimated polygenic score, we computed the PGS-corrected estimate of the exposure effect for ten replicates. We reported the mean of the estimated exposure effect across all ten replicates.

### Simulating genotypes and phenotypes

#### Random mating

Given that our empirical analyses rely on GWAS summary statistics obtained from the UK Biobank White British cohort, we conducted the majority of simulation analyses in simulated CEU individuals, except when analyzing portability (see *Analyzing portability* below). We simulated genotypes on chromosome 22 for CEU individuals under a published demographic model using the msprime engine in stdpopsim (Adrion *et al*., 2020; Baumdicker *et al*., 2022; Gutenkunst *et al*., 2009). For downstream analyses, we centered but did not standardize genotypes. To compare the utility of GRM- and PENGUIN-corrected estimates of exposure effect across different sample sizes, we randomly partitioned the simulated individuals into a GWAS cohort of 100,000 individuals and a test cohort of 4,000 individuals.

We simulated genetic values for exposure and outcome under an additive genetic architecture. To simulate causal variants, we first filtered variants for biallelic sites with a minor allele frequency greater than 0.01 in the test cohort. Next, we randomly sampled 10,000 variants to be the causal variants. We simulated effect sizes for the *i*th variant using a bivariate normal distribution

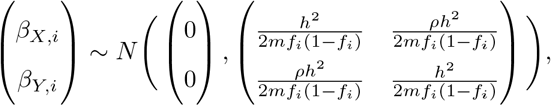

where *h*^2^ is the heritability of the exposure trait, *ρ* is the effect size correlation between exposure and outcome, *f*_*i*_ is the allele frequency of the *i*th variant in the test cohort, and *m* is the number of causal variants. This distribution corresponds to the assumption that each causal variant contributes equally to heritability (often known in statistical genetics as “the alpha model” (Speed *et al*., 2017)). For a subset of results, we also analyzed alternative models with a weaker dependence between effect size and frequency, such that Var(*β*_*i*_) ∝ (*f*_*i*_(1 − *f*_*i*_))^*α*^ for *α* ∈ [0, −0.3, −0.5].

We simulated trait-specific environmental effects as *N* (0, 1 − *h*^2^). Conditional on genetic values and environmental effects, we simulated phenotypes for exposure and outcome under the generative model we introduce in the Results:

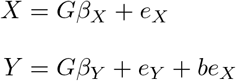

Note that this model ensures that both the variance of genetic values and the heritability of the exposure, *X*, is equal to *h*^2^, while only the variance of genetic values for the outcome is equal to *h*^2^; the heritability will vary with the magnitude of *b*.

#### Population structure

To model population structure, we simulated a population split followed by no migration between subpopulations. Specifically, we simulated a split time of 200 generations ago, and a constant population size of 10,000 individuals for each subpopulation. For downstream analysis, we sampled 52,000 individuals from each subpopulation. We simulated both stratified and unstratified phenotypes, and stratified phenotypes were simulated such that the environmental effect of both exposure and outcome was shifted by 0.5 units.

To compute GRM-corrected estimates, we analyzed a joint sample of 2,000 individuals from each subpopulation. To compute PENGUIN-corrected estimates, we used a joint sample of 50,000 individuals from each subpopulation to perform a GWAS. We then used a (non-overlapping) joint sample of 2,000 individuals from each subpopulation to comprise individual-level data for PENGUIN. We computed LD scores in a random subset of 4,000 individuals from one population, and incorporated principal components as covariates to correct population structure for both the GWAS and PENGUIN itself.

#### Assortative mating

We simulated genotypes and phenotypes under assortative mating using the software package xftsim (Border *et al*., 2024). Given the computational complexity of assortative mating simulations, we were unable to simulate haplotypes at a sample size capable of computing PENGUIN-corrected estimates. To evaluate GRM-corrected estimates in simulations of assortative mating, we simulated founder haplotypes of 4,000 CEU individuals on chromosome 22 using stdpopsim (Adrion *et al*., 2020). We seeded our simulations with these founder haplotypes and simulated 5 generations of cross-trait assortative mating with correlations ranging from −0.6 to +0.6. We simulated causal variants, effect sizes, and phenotypes under the models described previously (see *Random mating* above for more details).

### Analyzing portability

#### Portability of PENGUIN-corrected estimates

To characterize the portability of PENGUIN-corrected estimates, we simulated 10,000 CEU, CHB, and YRI individuals under the demographic model inferred by Gutenkunst *et al*. (2009). We randomly sampled 10,000 causal variants from the pool of biallelic sites that were segregating at a frequency of 1 *×* 10^−4^ in all populations. We simulated multivariate effect sizes independent of frequency (i.e. with *α* = 0), given the impossibility of simulating frequency-dependent effects in a multi-population model. We simulated a genetic correlation of 0.25 between exposure and outcome and an exposure heritability of 0.5 in CEU. Using simulated frequencies and effect sizes, we computed the analytical expectation of the PENGUIN estimator as follows, where *β*_*X,k*_ and *β*_*Y,k*_ are the effect of the *k*th variant on the exposure *X* and the outcome *Y*, and *p*_*k*,ind_ and *p*_*k*,GWAS_ are the frequency of the *k*th variant in the individual-level and GWAS cohorts respectively. (For a full derivation, see Supplementary Note 2.)

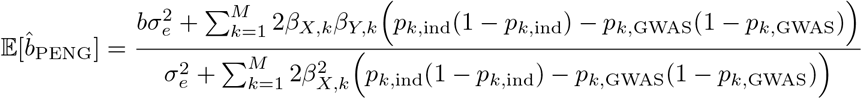

Across each of 10 simulation replicates, we computed the squared bias as 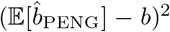 and reported the mean squared bias across simulations.

#### Portability of PGS-corrected estimates

To characterize the portability of PGS-corrected estimates, we simulated 100,000 CEU and 10,000 CHB and YRI individuals under the demographic model inferred by Gutenkunst *et al*. (2009). We simulated causal variants and their effect sizes as described in *Portability of PENGUIN-corrected estimates*. We generated GWAS summary statistics with plink2 (Chang *et al*., 2015) in the set of variants with MAF ≥ 0.01 in CEU, and clumped variants with *p* ≤ 1 *×* 10^−5^, *r*^2^ ≤ 0.5, and a window size of 250 kb. We computed PGS by summing over the product of clumped variants and their estimated effect sizes, and reported the mean squared error in each population.

### UK Biobank analyses

We analyzed the effect of loneliness on three cardiovascular traits (BMI, HDL cholesterol, and LDL cholesterol); four mental health traits (frequency of depressed mood, unenthusiasm, tiredness, and tenseness); and two immune-related traits (C-reactive protein and white blood cell count). To encode loneliness phenotypes, we used responses to the question “Do you often feel lonely?”, where “No” was encoded as 0, and “Yes” was encoded as 1. To encode categorical mood frequency phenotypes, we used the default encoding provided by the UK Biobank, where 1 corresponds to “Not at all”, 2 corresponds to “Several days”, 3 corresponds to “More than half the days”, and 4 corresponds to “Nearly every day”. To increase the interpretability of exposure effect, we inverse normal transformed phenotypes for quantitative traits (white blood cell count, C-reactive protein, LDL cholesterol, HDL cholesterol, and BMI) and removed outliers with a z-score greater than 3. We separately computed GRM- and PENGUIN-corrected estimates in three subsets of the UK Biobank: individuals who self-identified their ethnic background as Black or Black British (hereforth “Black British”); Indian; and both White and British (hereforth “White British”).

To compute GRM-corrected estimates, we constructed a GRM in each subset using all genotyped variants with minor allele frequency greater than 0.01. Given that there were 442,719 individuals who identified as White British, we randomly sampled 7,000 individuals to obtain a tractable sample size for computing GRM-corrected estimates. To compute GRM-corrected estimates in the Black British and Indian subsets, we used data from all 8,483 Black British and 6,104 Indian individuals. We performed variance components analysis using bivariate GREML and included covariates that captured age, sex, assessment center, and 10 principal components.

We computed PENGUIN-corrected estimates using GWAS summary statistics obtained from all UK Biobank White British individuals (http://www.nealelab.is/uk-biobank/). We used individual-level phenotype data from the random sample of 7,000 White British individuals and all Black British and Indian individuals. As before, we included covariates that captured age, sex, and 10 principal components.

## Data and code availability

Software for generating GRM-corrected estimates of exposure effect and code for replicating results in this manuscript is available at https://github.com/roshnipatel/grm-corrected-exposure.

## Acknowledgments and funding sources

We thank members of the Edge, Pennell, and Mooney labs for helpful conversations about this work. We are also grateful to Graham Coop for thoughtful comments on an earlier draft. RAP, JGS, and MDE were supported by NIH grant R35GM137758. MP was supported by NIH grant R35GM151348.

## Supplementary Note 1

We will show that the generative statistical model described in the main text can be understood as underlying existing methods to control genetic confounding in observational epidemiological studies. That is, under favorable conditions for each method, existing methods estimate the same parameter as our method under the model we assume. As in the main text, suppose we have two traits *X* and *Y*, such that:

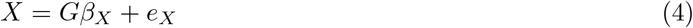

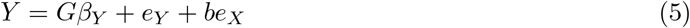

where *G* is the centered genotype matrix; *β*_*X*_ and *β*_*Y*_ are the additive SNP effects; and *e*_*X*_ and *e*_*Y*_ are the non-genetic effects on the traits. We will refer to *X* as the exposure and *Y* as the outcome. We assume *e*_*X*_ and *e*_*Y*_ are independently and identically distributed across individuals with a mean of 0 and that *e*_*X*_ and *e*_*Y*_ are uncorrelated with each other. (The non-genetic components of *X* and *Y* can still be correlated, as the non-genetic component of *Y* is *e*_*Y*_ + *be*_*X*_.) We further assume that non-genetic effects *e*_*X*_ and *e*_*Y*_ are independent of *G*. We assume *β*_*X*_ and *β*_*Y*_ have a mean of 0 and that *β*_*X*_ and *β*_*Y*_ could be independent or correlated.

Under this model, the desired estimate of exposure effect is 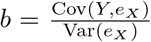. Regressing *Y* on *e*_*X*_ is generally not possible because *e*_*X*_ is unmeasurable. Thus, the naive approach is to regress *Y* on *X*. In expectation, the estimate of the exposure effect obtained from this naive regression is:

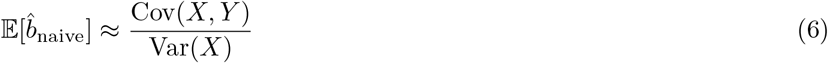

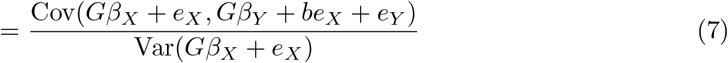

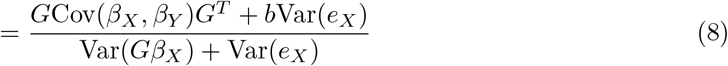

It is evident that the naive estimate of exposure effect is subject to bias from the genetic correlation between exposure and outcome, Cov(*β*_*X*_, *β*_*Y*_). However, even when this term is 0 (i.e. if there is no genetic correlation between traits), the expectation of the naive estimate of exposure effect simplifies to:

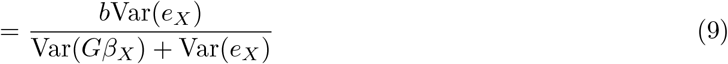

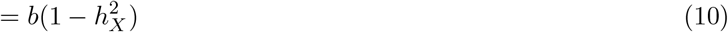

In other words, the naive estimate of exposure effect is *also* subject to regression attenuation due to the noise generated by unmodeled genetic variation in the exposure. Thus, the naive estimator is not appropriate for estimating *b*, the interventional effect of the exposure.

We will next show that this is the same model underlying PGS-based methods to control genetic confounding. In other words, we will show that PGS-corrected estimates of the exposure effect seek to estimate *b* as defined above. If we suppose *X* is a function of additive genetic effects, *Gβ*_*X*_, and non-genetic effects, *e*_*X*_, a perfect PGS for *X* is given by *Gβ*_*X*_. PGS-based methods for controlling genetic confounding include the exposure PGS as a covariate in the regression of outcome on exposure:

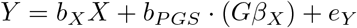

In expectation, the estimate of the coefficient *b*_*X*_ is:

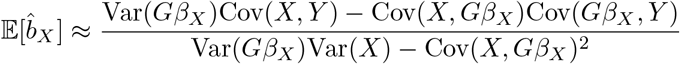

where the approximation results from approximating the expectation of a random quotient by the quotient of the expectations of its random numerator and denominator. Assuming independence of genetic and non-genetic effects,

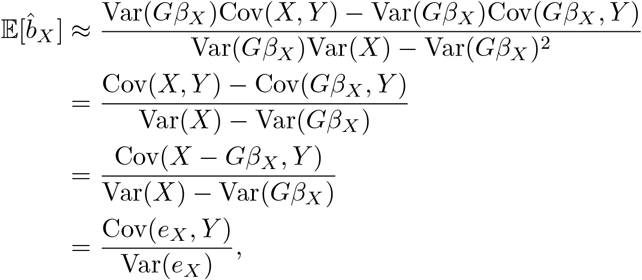

and we have again recovered the interventional effect of the exposure, as defined in Equation 10. Thus, PGS-corrected estimates are equivalent to *b*, the interventional effect of the exposure when the exposure PGS is identical to the true genetic value. However, we note that this is not true in the absence of the perfect PGS for the exposure.

Specifically, we wish to note that regressing on the perfect PGS for the outcome does not control genetic confounding, as previously suggested (Zhao *et al*., 2024). In particular, using the perfect

PGS for the outcome yields the following estimator:

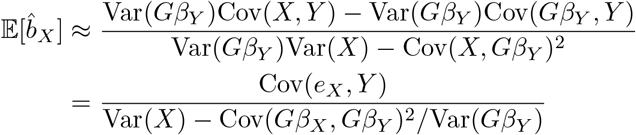

By the Cauchy–Schwarz inequality, regressing on the perfect PGS for the outcome will only yield the correct estimate if *β*_*Y*_ is collinear with *β*_*X*_.

Finally, we will show that PENGUIN assumes the same generative model and seeks to approximate the same parameter (Zhao *et al*., 2024). The PENGUIN estimator is introduced as:

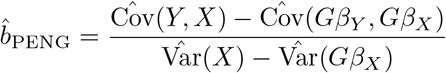

Then, in expectation, the PENGUIN estimator will approximately be a function of the true variances and covariances. Using the independence of *β*_*X*_, *β*_*Y*_, *e*_*X*_, and *e*_*Y*_,

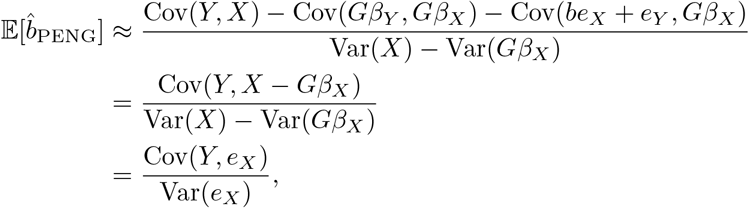

and we have thus recovered the interventional effect of the exposure, as defined in Equation 6.

## Supplementary Note 2

We will evaluate the portability of PENGUIN-corrected estimates by deriving an analytical expression for the PENGUIN estimator when GWAS summary statistics are obtained from one cohort and individual-level data is obtained from a second cohort.

As before, suppose we have an exposure *X* and an outcome *Y* influenced by genetic factors *G* and non-genetic factors *e*_*X*_ and *e*_*Y*_, and let *G*_GWAS_ and *G*_ind_ correspond to the genotype matrix in the GWAS and individual-level cohorts, respectively.

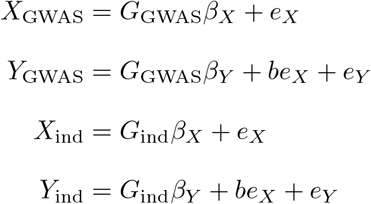

We will assume that the non-genetic effect of the exposure, *b*, is constant across both cohorts in absolute units. We will also assume that the total variance contributed by non-genetic factors (i.e. Var(*e*_*X*_) and Var(*e*_*Y*_) is shared across both cohorts. (Note, of course, that in real-world situations, environmental variance can differ among groups (Mostafavi *et al*., 2020; Wang *et al*., 2024).)

In a scenario where the PENGUIN estimator is constructed from two different cohorts, the estimator is:

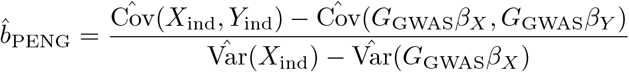

In expectation, 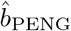 is not necessarily equal to *b*. If we take the expectation of 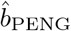 over *G, e*_*X*_, and *e*_*Y*_, and assuming independence of *G, e*_*X*_, and *e*_*Y*_, we have:

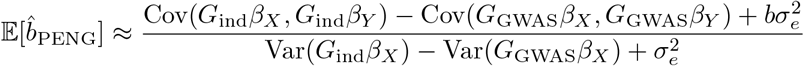

If we assume that genotypes at causal loci are independent within each cohort, we can simplify further. Let *β*_*X,k*_ and *β*_*Y,k*_ be the effect of the *k*th variant on the exposure *X* and the outcome *Y*, and let *p*_*k*,ind_ and *p*_*k*,GWAS_ be the frequency of the *k*th variant in the individual-level and GWAS cohorts respectively.

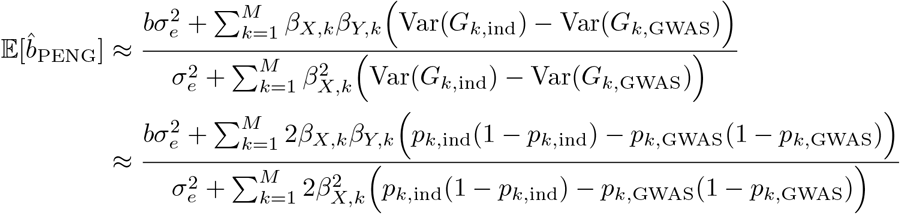

It should be fairly straightforward that, if *p*_*k*,GWAS_ and *p*_*k*,ind_ are identical, 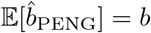. Alternately, if *p*_*k*,GWAS_ and *p*_*k*,ind_ are not identical, and per-SNP heritabilities differ between the two cohorts, 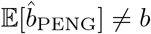 except in special cases.

## Supplementary Figures

**Figure S1.**
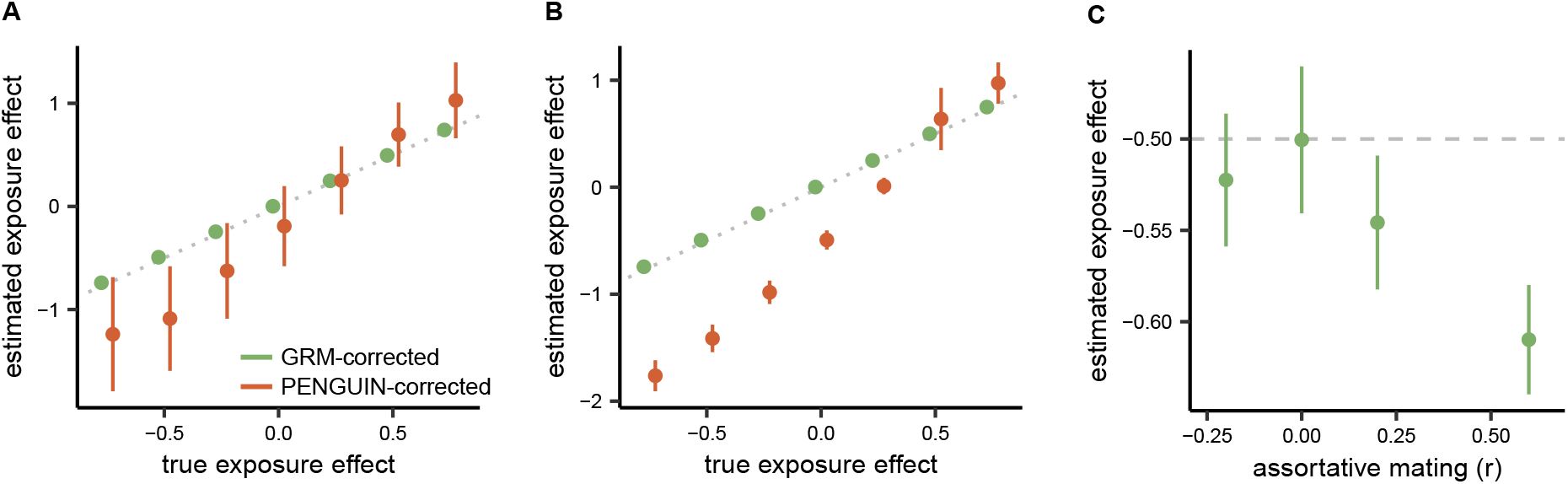
Non-random mating. **A)** GRM- and PENGUIN-corrected estimates of exposure effect in simulations of population structure, but no mean differences in environmental effect. **B)** GRM- and PENGUIN-corrected estimates of exposure effect in simulations of population stratification, i.e. a mean difference of 0.5 units in the environmental component for both exposure and outcome. **C)** GRM-corrected estimates of exposure effect across simulations of assortative mating; the true exposure effect is simulated at −0.5. Across all three simulations, exposure heritabilities were simulated at 0.5 in the absence of assortment and structure. Error bars correspond to mean *±* 1 standard deviation.

**Figure S2.**
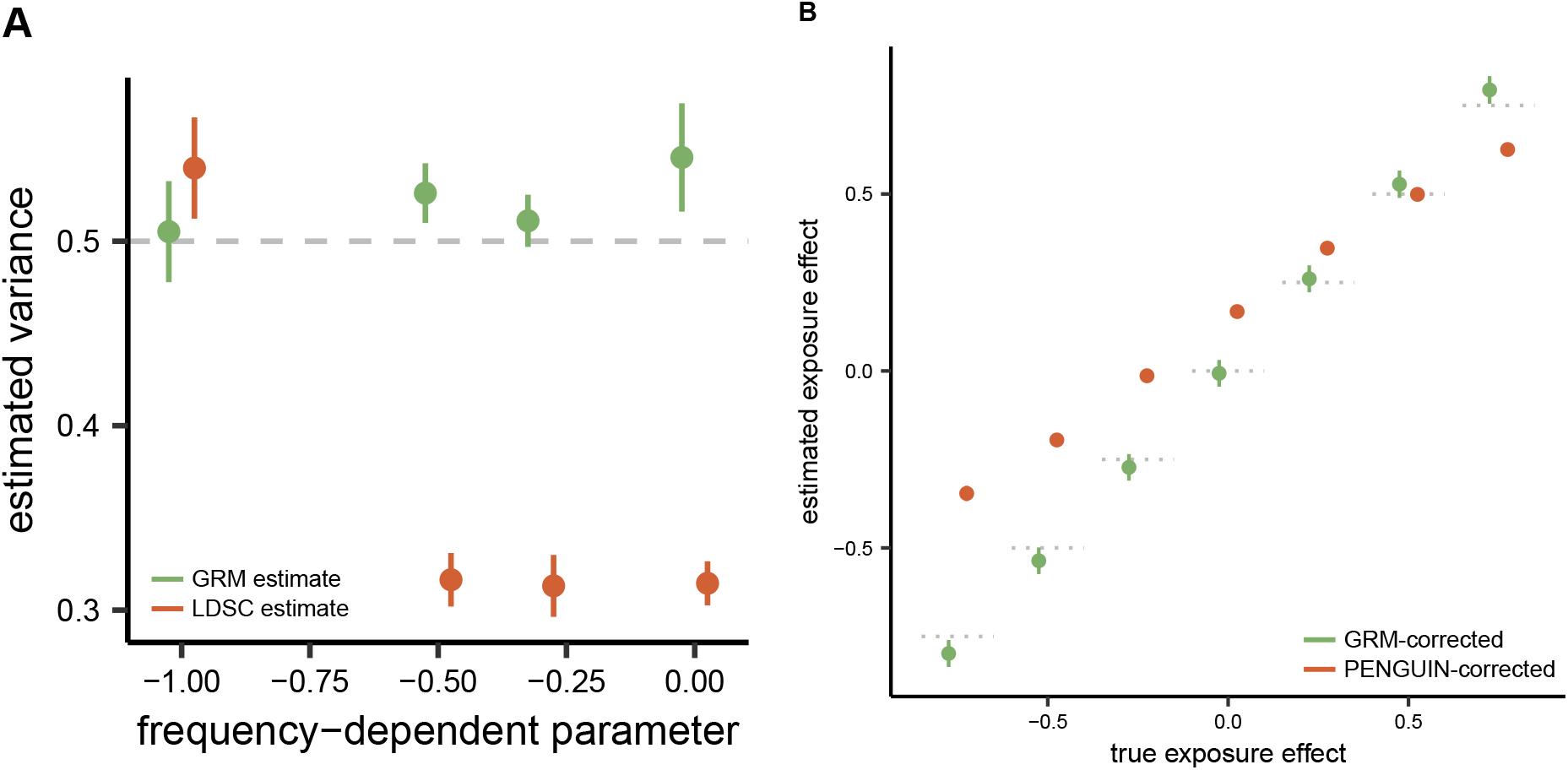
Heritability models. **A)** Estimates of heritability from GRMs and LDSC across simulations with different relationships between allele frequency and effect size. The x-axis corresponds to the parameter *α* (see Methods), such that *α* = − 1 represents the default assumption of the canonical GRM and of PENGUIN, and *α* = 0 represents frequency-independent effects. The true simulated heritability is 0.5. **B)** GRM- and PENGUIN-corrected estimates of exposure effect when *α* = 0 and allelic effects are independent of frequency. Horizontal dotted lines correspond to the diagonal, i.e. the *y* = *x* line. Error bars correspond to mean *±* 1 standard deviation.

**Figure S3.**
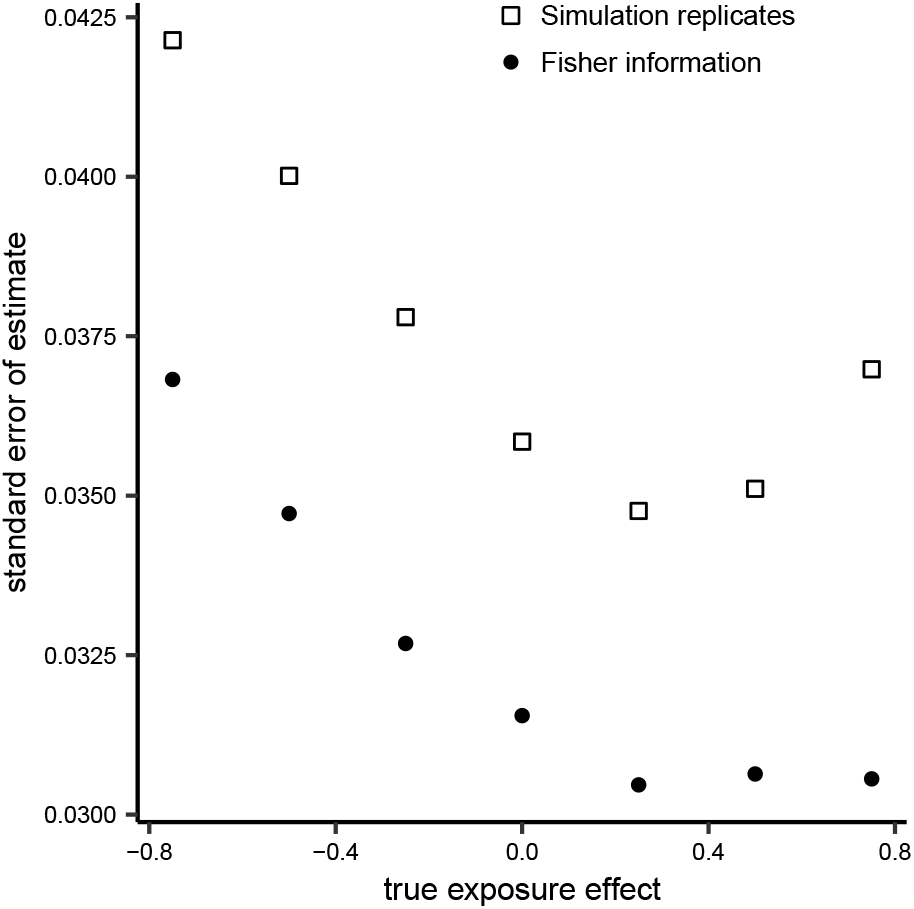
Comparison of standard errors. Standard errors of GRM-corrected estimates obtained from 10 simulation replicates compared with standard errors computed via the Fisher information matrix associated with a single simulation replicate.

## Notes

### Competing Interest Statement

The authors have declared no competing interest.

### Summary of Updates

Discussion and Supplementary Note 1 revised; Figure 5 added; more details on theory.

